# The mistletoe *Struthanthus flexicaulis* reduces dominance and increases diversity of plants in campo rupestre

**DOI:** 10.1101/2020.04.30.070268

**Authors:** Graziella França Monteiro, Milton Barbosa, Yasmine Antonini, Marcela Fortes de Oliveira Passos, Samuel Novais, G. Wilson Fernandes

## Abstract

The interaction between hemiparasites and their host plants is an important structuring mechanism for plant communities. The mistletoe *Struthanthus flexicaulis* (Loranthaceae) is widely distributed in the campo rupestre ecosystem and likely has an important role in structuring the communities of which its hosts are part. The main goals of this study were to investigate the effects of parasitism by *S. flexicaulis* on host plants in a degraded area of campo rupestre and to determine how parasitism affects characteristics of the structure of this plant community over time. We found that parasitized plants had smaller crowns and branch growth, and suffered lower mortality compared to non-parasitized plants. Parasitism by *S. flexicaulis* decreased dominance and increased the diversity and evenness of plants in the community over time. Parasitism leads to competition with the host for water and nutrients, which may decrease the performance of the host and, consequently, leading to host death. The high mortality of the most abundant plant species led to a restructured woody plant community. These results reinforce the importance of parasitic plants as key species for maintaining species diversity in plant communities.

## INTRODUCTION

Identifying the factors that control the distribution, abundance and diversity of species is important to understand the structuring of ecological communities. Communities are structured according to climatic and historical variations (e.g. dispersion and speciation, migration and extinction), local physical conditions and interactions among species [1, 2]. On a local scale, abiotic factors have an important influence on the structuring of plant communities, such as the availability of water and soil nutrients [34] in addition to interactions with other organisms, such as microbes and arbuscular mycorrhizal fungi [5, 6, 7]. In turn, the distribution of plant communities strongly affects the community structure of other organisms that directly depend on them as resources, such as herbivores and parasitic plants [8, 9, 10].

Among parasitic plants, hemiparasites are those capable of photosynthesis, but depend on their hosts for water and nutrients [11]. The susceptibility to attack by hemiparasitic plants (determined by chemical, physiological and physical processes at the parasite-host interface) varies between plant species, and therefore, local abundance and the degree of susceptibility of plants in a community are key factors for the successful colonization of hemiparasites in the environment [12, 13, 14]. In addition, the seeds of the hemiparasitic plants are mostly dispersed by birds, which can lead to a differential deposition of seeds on the host plants according to the preferences of the birds for certain perches [15, 16, 17]. Generally, larger plants with more branched tops are more attractive perches for birds, leading to a greater abundance of hemiparasitic plants on these hosts [15, 18, 16, 19, 20].

Once the hemiparasitic plants are established, they can affect the host plant both directly and indirectly [21, 22, 23, 9] and, consequently, affect the structure of plant communities [14]. Directly, the hemiparasite decreases the growth and branching of the host plant [23], as they obtain water and nutrients directly from the xylem, decreasing the resources available for the metabolism of the host itself [11, 24, 25], which may ultimately lead to death [19, 26, 9]. Indirectly, the parasite can make the host plant more susceptible to attack by natural enemies [23] and to environmental stress [12, 11], which can also accelerate the death of the host. Therefore, once the most abundant plant species in the environment are more frequently parasitized, the hemiparasitic plant can play an important role in the structuring of plant communities, since by promoting a higher mortality of individuals of dominant species, it would lead to greater uniformity in the number of individuals among plant species in the community [11, 27, 28, 29, 30, 31].

Loranthaceae is one of the most representative families of parasitic plants in Brazil, with 10 genera and approximately 100 species described [32, 14]. In the Cerrado biome, 36 species have been described, all of which are hemiparasites [33]. Among them, the mistletoe *Struthanthus flexicaulis* is a common generalist hemiparasite in campos rupestres, with more than 60 species recorded as hosts within at least 24 families of woody plants [25, 9]. The campo rupestre ecosystem is subject to significant anthropic pressure such as mining and road construction [3, 34], and in degraded areas the native pioneer shrub, *Baccharis dracunculifolia,* whose populations form patches, often dominates the plant community [3, 10].

Casual observations indicated a high incidence of the mistletoe S. *flexicaulis* on the patches of *B. dracunculifolia*, therefore representing an excellent experimental model to test whether the plant community can be shaped by the presence of a plant parasite. This study aimed to evaluate the factors related to the host plants that affect the distribution and abundance of this important and widespread mistletoe, and the effects of hemiparasitism on host plants in a degraded area of campo rupestre, in southeastern Brazil. We tested whether the presence of the S. *flexicaulis* affects the structure of this community over time. First, we determined the effect of the frequency and architecture of the woody host plants on parasitism by *S. flexicaulis*. Predictions of the following hypotheses were tested: (1) More common plant species in the environment are more attacked by the *S. flexicaulis*; (i) more abundant host plants would present a greater intensity of infection (number of parasites per host) by *S. flexicaulis*; (2) Structurally more complex host plants would be more parasitized by *S. flexicaulis*; (i) host plants with more complex architecture (plant height and crown, number of primary and secondary branches and maximum branching) would present a higher intensity of infection by *S. flexicaulis*; (ii) Branches in greater diameter classes would be more parasitized by *S. flexicaulis* compared with smaller diameter classes. In a second step, we determined the impact of parasitism by *S*. *flexicaulis* on host plants and the woody plant species community. The predictions of the following hypotheses were tested: (1) Parasitized plants show lower growth and suffered higher mortality than non-parasitized plants: (i) parasitized plants would show lower growth (plant height and crown, number of primary and secondary branches, and maximum branching) over time compared to non-parasitized plants; parasitized plants would have a higher proportion of individuals killed over time compared to non-parasitized plants; (2) Parasitic plants favor the diversity of woody plants, reducing the abundance of dominant species over time: (i) the dominance of species in the woody plant community with the presence of the mistletoe *S. flexicaulis* would decrease over time, while diversity and evenness would increase.

## MATERIALS AND METHODS

### Study area

The study was conducted at Reserva Vellozia (19°16’45.7” S 43°35’27.8”W), in Serra do Cipó, southern portion of the Espinhaço Range, in Minas Gerais, Brazil. Reserva Vellozia is located at an altitude between 900-1355 meters, where the vegetation of campos rupestres is predominant. The vegetation of campo rupestre in Serra do Cipó is characterized by a high richness of plant species, with about 125 families of phanerogams with more than 2000 species of plants listed [35]. In addition, it presents high levels of endemism and several endangered species [3, 36, 34, 37]. The soil is shallow and often prevents large tree species from being established [38].

### Sample design

In March 2017, eight 2 m x 50 m transects were established over 4 km, at least 500 m apart. The transects were located in areas under anthropic pressure due to the construction of roads, with a high incidence of the mistletoe *S. flexicaulis*. In each transect, all woody plants (parasitized and non-parasitized) taller than one meter were marked (plants shorter than one meter are generally not parasitized; [23] and the richness (number of species) and abundance (number of individuals) were recorded. Specimens of all the plant species marked were identified to the lowest possible taxonomic level. In addition, the number of *S*. *flexicaulis* individuals established (with green leaves and fixed on the stem of the host plant) on all parasitized hosts was recorded.

The following measurements of the architecture of all marked plants (parasitized and non-parasitized) were taken (using a measuring tape): height of the plant (from the ground to the top of the crown), height of the crown (first branch to the top), number of primary branches, number of secondary branches and maximum level of branching (maximum number of branching levels that a plant has; methodology adapted from [39]. We measured the diameter (Mitutoyo caliper: mm) of each parasitized branch, which was grouped according to five diameter classes (0-2 mm, 3-5 mm, 6-9 mm, 9-15 mm and >15 mm). To compare mortality and growth between parasitized and non-parasitized plants, we performed a new measurement, in July 2019 (28 months after the first one), of the same architectural parameters (height, number of primary branches, number of secondary branches, maximum branching) and we quantified the living and dead plants. The proportion of dead parasitized and non-parasitized individuals per transect was calculated. The growth of each individual was calculated by the difference in plant height and crown height between the years 2017 and 2019. The same was done for the increase in the number of primary branches, secondary branches and maximum branching, as they were calculated by the difference between the measurements of the years 2017 and 2019.

### Data analysis

To assess whether the most frequent host plant species in the environment show a higher intensity of infection (number of parasites per host) by *S. flexicaulis,* a mixed generalized linear model (GLMM) [40] was used. The response variable was the average number of individuals of *S. flexicaulis* per individual of each species of host plant per transect, the fixed variable was the frequency of each host plant species per transect and the random variable was the sampled transects.

A GLMM was used to assess whether individuals from structurally more complex host plants have a higher intensity of *S*. *flexicaulis* infection. The response variable was the total number of individuals of *S. flexicaulis* per individual of the host plant, the fixed variables were height, crown height, number of primary and secondary branches and maximum number of branches of the individuals of the host plants, and the random variable was the sampled transects. To observe if branches with larger diameters are more parasitized, a GLMM was created, where the response variable was the abundance of the parasitic plants per transect, the fixed variable was the size classes and the random variable was the transects. To evaluate the mortality of the host plants, a GLMM was performed, where the response variable was mortality after 28 months, the fixed variable was the presence or absence of the mistletoe *S. flexicaulis* in the plants and the random variable was the sampled transects. The variables of plant height and plant crown were used as covariates to observe the effect on mortality. To evaluate whether the presence of the mistletoe *S. flexicaulis* affects the growth of host plants after 28 months, GLMMs were performed. The growth measures were the response variables of each model, the fixed variable was the presence or absence of the mistletoe *S. flexicaulis*, and the random variable was the transects. We performed all analyzes in the R Software [41]. Finally, to determine if there are differences in the structure of the plant community with the presence of the mistletoe *S. flexicaulis* between the years 2017 and 2019, we used Simpson’s dominance index, Pielou's evenness index, and Shannon’s diversity index [42]. As the objective of evaluating how the structure of the plant community would behave over time without the presence of the parasite, the same indexes were calculated separately for the group of non-parasitized plants between the years 2017 and 2019.

## RESULTS

In the first year of study (2017) 271 individuals belonging to 11 families and 20 species of plants were recorded in the eight sampled transects. The most representative families in number of species were Asteraceae (four species) and Fabaceae (three species; Table 1). Among all sampled species, *Baccharis dracunculifolia* was the most abundant (n = 189), followed by *Callisthene* sp (n = 17) and *Mimosa maguirei* (n = 11; Table 1). Among the 271 individuals and 20 recorded species, 151 individuals (55%) and nine species (45%) were parasitized by *S. flexicaulis*. The greatest abundance of individuals from *S. flexicaulis* parasitizing a single host plant (n = 9) was found in *B. dracunculifolia*. Less frequent species, such as *M. maguirei* (n = 11) and *Solanum lycocarpum* (n = 6) were not parasitized (Table 1).

**Table 1.** Plant community parasitized and not parasitized by *Struthanthus flexicaulis* in a disturbed area of campos rupestres, in Serra do Cipó MG, Brazil. Host abundance: the total abundance of each species of parasitized and non-parasitized plants. Mistletoe abundance: the abundance of parasites by plant species. % Infected: the proportion of individuals of each species infected with at least one individual of *S. flexicaulis*. Host mortality: the abundance and percentage of dead plants.

| Family | Species | Host<br>abundance | Mistletoe<br>abundance | %<br>Infected | Host<br>mortality |
| --- | --- | --- | --- | --- | --- |
| Asteraceae | *Baccharis dracunculifolia* | 189 | 132 | 0,67 | 117 - 0,61 |
| Asteraceae | *Eremanthus erythropappus* | 1 | 1 | 1,00 | 0 - 0,0 |
| Asteraceae | *Eremanthus incantus* | 2 | 0 | 0,00 | 0 - 0,0 |
| Asteraceae | *Vernonanthura scorpioides* | 2 | 0 | 0,00 | 0 - 0,0 |
| Clusiaceae | *Kielmeyera petiolaris* | 5 | 1 | 0,25 | 2 - 0,50 |
| Clusiaceae | *Kielmeyera scorpioides* | 5 | 0 | 0,00 | 2 - 0,40 |
| Fabaceae | *Stryphnodendron adstringens* | 1 | 0 | 0,00 | 0 - 0,0 |
| Fabaceae | *Dalbergia miscolobium* | 4 | 0 | 0,00 | 0 - 0,0 |
| Fabaceae | *Mimosa maguirei* | 11 | 0 | 0,00 | 0 - 0,0 |
| Lamiaceae | *Hyptidendron asperrimum* | 2 | 3 | 1,0 | 2 - 1,0 |
| Lamiaceae | *Hyptidendron canum* | 1 | 0 | 0,00 | 0 - 0,0 |
| Lythraceae | *Diplusodon orbicularis* | 8 | 3 | 0,25 | 4 - 0,50 |
| Malpighiaceae | *Byrsonima verbascifolia* | 1 | 0 | 0,00 | 0 - 0,0 |
| Malpighiaceae | *Byrsonima* | 8 | 2 | 0,25 | 3 - 0,37 |
| Myrtaceae | *Psidium firmum* | 1 | 0 | 0,00 | 1 - 1,0 |
| Salicaceae | *Casearia sylvestris* | 3 | 0 | 0,00 | 1 - 0,33 |
| Solanaceae | *Solanum lycocarpum* | 6 | 0 | 0,00 | 0 - 0,0 |
| Urticaceae | *Cecropia pachystachya* | 1 | 1 | 1,0 | 0 - 0,0 |
| Vochysiaceae | *Callisthene* | 17 | 7 | 0,35 | 7 - 0,41 |
| Vochysiaceae | *Vochysia thyrsoidea* | 3 | 4 | 0,66 | 1 - 0,33 |

The host species that were more frequent in the environment showed a greater abundance of parasitic plants per individual (intensity of infection; Tables 1 and 2; Fig 1a). Positive associations between the abundance of S. *flexicaulis* and the height of the plants (Fig 1b), the height of the crowns (Figure 1c) and the maximum level of branching (Table 2; Fig 1d) were recorded. On the other hand, the number of primary and secondary branches of the plants did not affect the abundance of *S. flexicaulis* per plant (Table 2). In addition, the abundance of *S. flexicaulis* was higher in the diameter classes 2 and 3 (Table 2; Fig 2a).

**Table 2.**
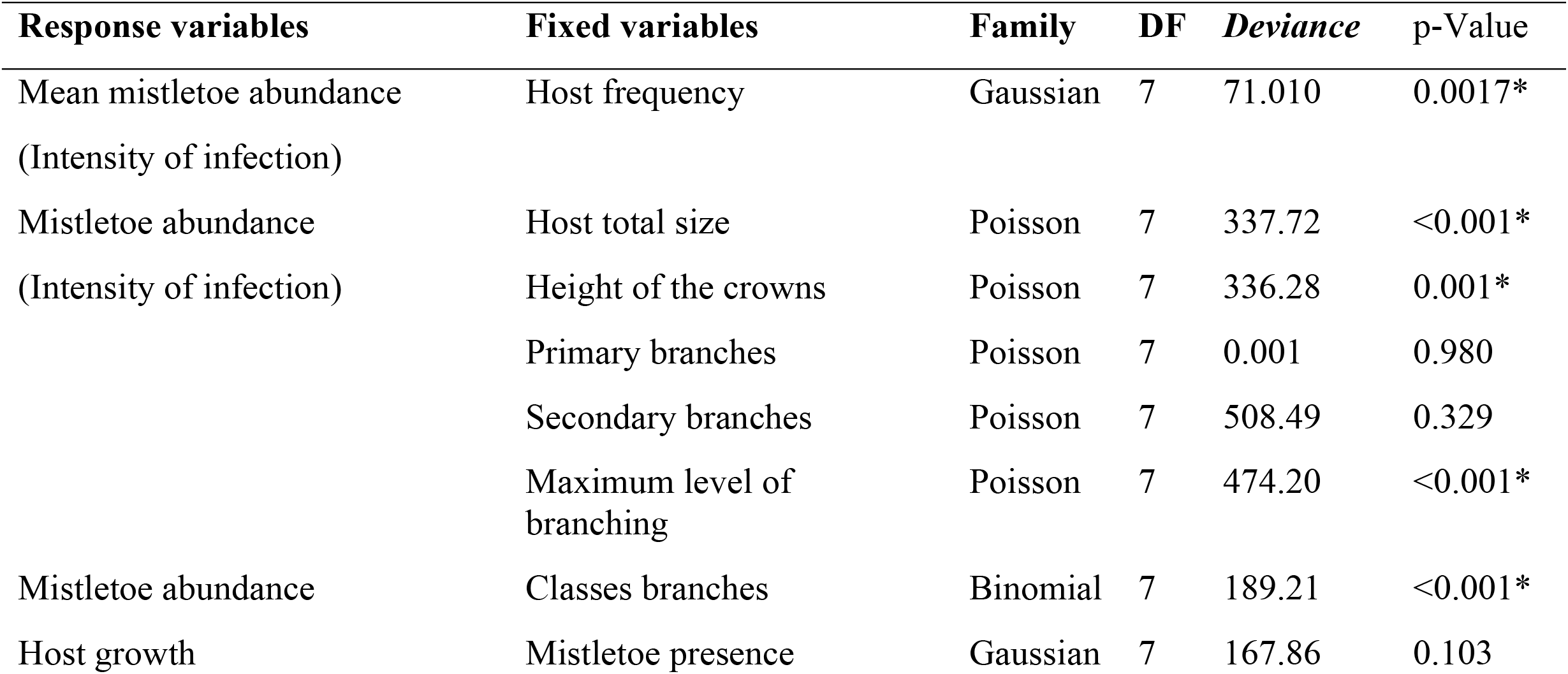
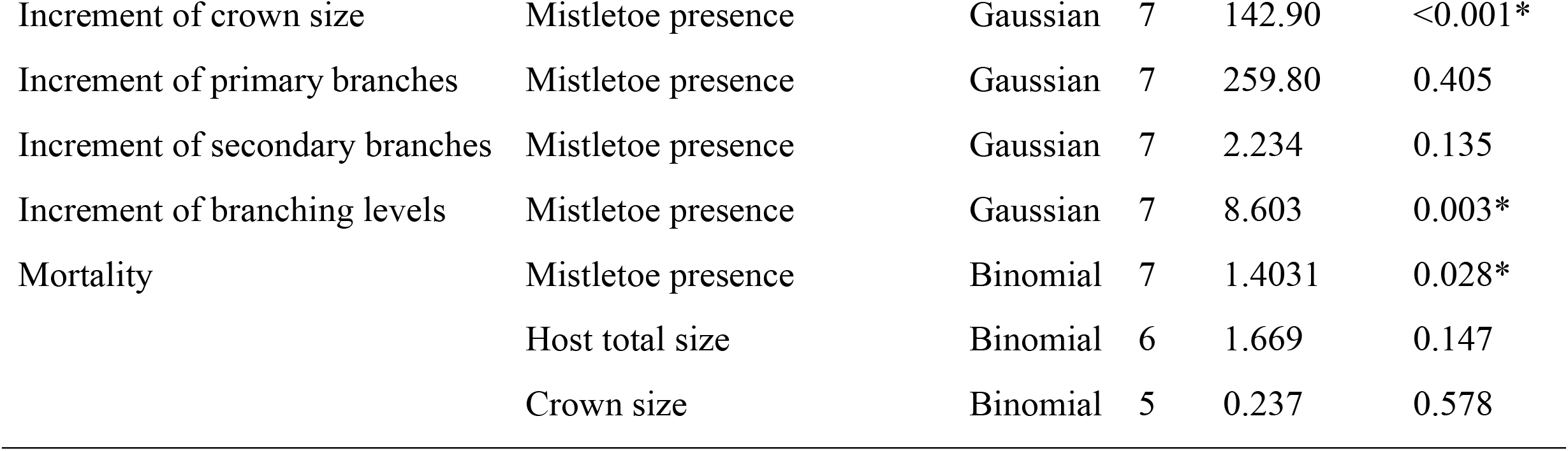
Results of generalized linear mixed models (GLMM) testing the effect of host plant frequency and architecture on the abundance of the mistletoe *Struthanthus flexicaulis*, and the effect of this parasite on the growth and mortality of host plants in campos rupestres, in Serra do Cipó, MG, Brazil.

| Response variables | Fixed variables | Family | DF | Deviance | p-Value |
| --- | --- | --- | --- | --- | --- |
| Mean mistletoe abundance<br>(Intensity of infection) | Host frequency | Gaussian | 7 | 71.010 | 0.0017* |
| Mistletoe abundance<br>(Intensity of infection) | Host total size | Poisson | 7 | 337.72 | <0.001* |
|  | Height of the crowns | Poisson | 7 | 336.28 | 0.001* |
|  | Primary branches | Poisson | 7 | 0.001 | 0.980 |
|  | Secondary branches | Poisson | 7 | 508.49 | 0.329 |
|  | Maximum level of<br>branching | Poisson | 7 | 474.20 | <0.001* |
| Mistletoe abundance | Classes branches | Binomial | 7 | 189.21 | <0.001* |
| Host growth | Mistletoe presence | Gaussian | 7 | 167.86 | 0.103 |
| Increment of crown size | Mistletoe presence | Gaussian | 7 | 142.90 | <0.001* |
| Increment of primary branches | Mistletoe presence | Gaussian | 7 | 259.80 | 0.405 |
| Increment of secondary branches | Mistletoe presence | Gaussian | 7 | 2.234 | 0.135 |
| Increment of branching levels | Mistletoe presence | Gaussian | 7 | 8.603 | 0.003* |
| Mortality | Mistletoe presence | Binomial | 7 | 1.4031 | 0.028* |
|  | Host total size | Binomial | 6 | 1.669 | 0.147 |
|  | Crown size | Binomial | 5 | 0.237 | 0.578 |

**Fig. 1.**
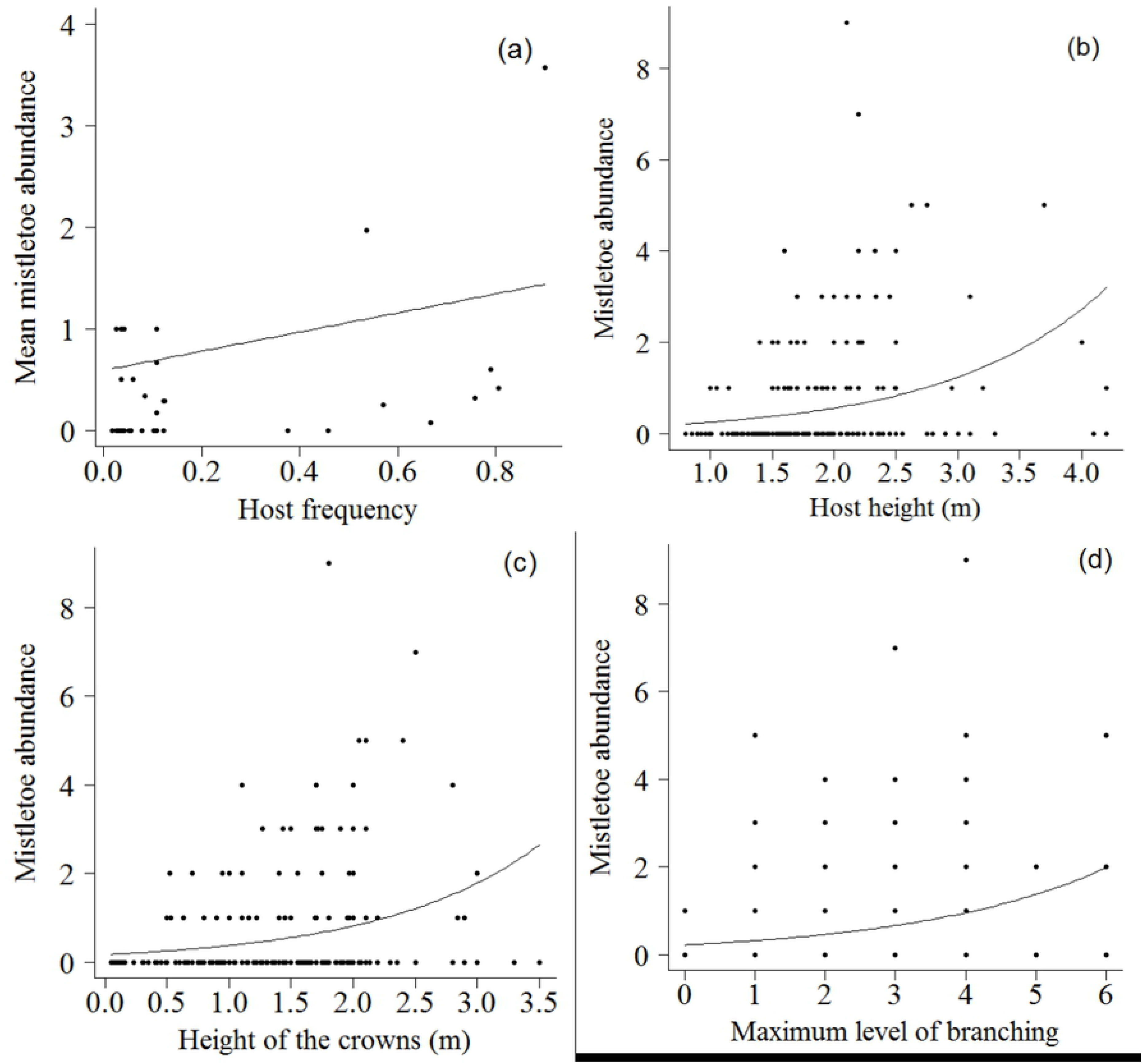
Relationship between intensity of infection (abundance of individuals per host plant) by mistletoe *Struthanthus flexicaulis* and (a) host frequency (y= −0.1348 + 33.3566*x), (b) host height (y= exp(−2.1663 + 0.7923*x)), (c) height of the crowns (y=exp (−1.7705 + 0.7827*x)), and (d) maximum level of branching (y=exp (−1.50740 + 0.36442*x)), in campos rupestres, in Serra do Cipó, MG, Brazil.

**Fig. 2.**
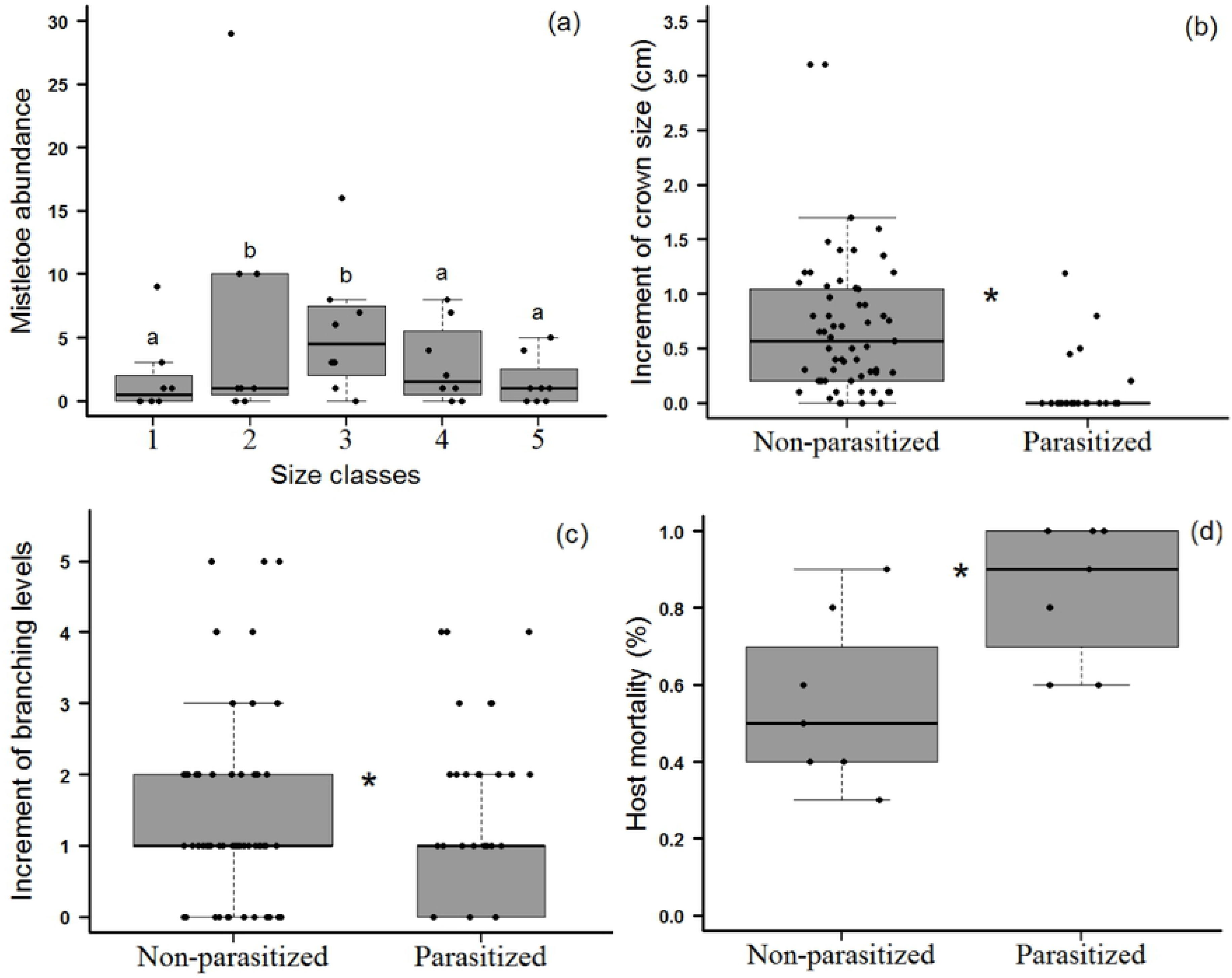
(a) Classes of parasitized branches, (b) increment of crown size, (c) increment of branching levels of plants (d) mortality of plants parasitized and non-parasitized and by the mistletoe *Struthanthus flexicaulis* in campos rupestres, in Serra do Cipó, MG, Brazil.

Parasitized plants showed lower crown growth and lower increment of branches over time than the non-parasitized plants (Table 2; Fig 2b and c). However, for the other architectural variables measured over time (height, primary, and secondary branches), no statistically significant differences were found between parasitized and non-parasitized plants (Table 2). Plant mortality was intensified by the presence of *S. flexicaulis* (Table 2; Figure 2d). Out of the 120 non-parasitized plants, 26 individuals (21.66%) died after 28 months, while among the 151 parasitized plants, 128 individuals (84.10%) died during the same period. Among the most abundant species (n > 7), *B. dracunculifolia* and *Diplusodon orbicularis* suffered higher mortality (Table 1). There was no effect of plant and canopy height on plant mortality (Table 2).

The community structure changed between the years of 2017 and 2019. There was a reduction in species richness from 20 to 18 species, since individuals of *Hyptidendron asperrimum* (n = 2) and *Psidium firmum* (n = 1) died. The plant community showed greater dominance in 2017 (Simpson = 0.495) than in 2019 (Simpson = 0.322). In addition, evenness was lower in 2017 (Pielou = 0.201) than in 2019 (Pielou = 0.274). Species diversity was lower in 2017 (Shannon = 0.603) compared to 2019 (Shannon = 0.793). When we evaluated only the non-parasitized plant group, we observed that the structure of this assemblage remained stable between the years 2017 (Simpson = 0.246; Pielou = 0.0537; Shannon = 0.860) and 2019 (Simpson = 0.242; Pielou = 0.0572; Shannon = 0.856).

## DISCUSSION

The abundance of the mistletoe *S. flexicaulis* depends directly on the abundance of host plants in the studied community, that is, the more frequent host species in the community were more intensely infected (number of hemiparasites per individual). In our study, the most frequent and most parasitized plant species, *Baccharis dracunculifolia*, is a pioneer plant that occurs in dense patches [43, 44, 45]. Then, these patches may become more visible and represent important resting sites to birds in disturbed areas, increasing the deposition of seeds and consequently the abundance of parasitic plants [46], [19]. In a study carried out in ferruginous campo rupestre, [9] found that *Mimosa calodendron* was the host plant with the highest proportion of parasitized individuals. The authors suggested that the aggregate populations of this host plant within the community, forming green patches in the landscape, determined its greater use as perches by seed-dispersing birds and, consequently, a greater deposition of hemiparasite seeds. Many individuals of the same species, in dense patches, can also favor the spread of the parasite, since part of the seeds ingested by the birds are left close to the mother plant [46], [47], [25]. In addition, the high density of host plants in patches can increase the likelihood of individuals receiving branches of the parasite from neighboring infected plants [9].

Plants with more complex architecture had more *S. flexicaulis*. Taller plants with larger and more branched crowns are more likely to be used as perches and offer a greater number of sites for the establishment of *S. flexicaulis* [19, 20, 23]. Greater abundance of *S. flexicaulis* on individuals of *B. dracunculifolia* that presented larger classes of size and with more levels of branching were reported by [23]. In addition, plants with more complex architecture have branches with a greater variety of sizes, which can increase the amount of optimal substrates for the establishment of the hemiparasite. Similarly, to our results, previous studies have shown that branches with intermediate diameters are the most colonized, as in them the parasite’s haustorium penetration is facilitated, while parasitic plants are unable to establish themselves in branches with very small diameters and very large branches have very thick bark that prevents haustorium penetration [48, 18]. Another important factor is that hosts with more complex architecture, probably older ones, have been in the environment for a longer time, and therefore, they were more likely to be infected.

Plants parasitized by *S. flexicaulis* showed lower crown growth and branching than non-parasitized plants, over time. This result was expected once the parasitic plant competes with the host plant for water and nutrients [49]. Other studies have shown that host plants attacked by *S. flexicaulis* showed lower foliage cover compared to non-parasitized plants [19, 23]. In addition to interfering with the vigor of parasitized plants, parasitism can affect the fitness of their hosts [49, 50]. [19] demonstrated that individuals of *M. calodendron* parasitized by *S. flexicaulis* produce less fruits and lighter seeds than those not parasitized, suggesting that the viability of these seeds can also be reduced. Particularly, in environments under water stress, such as the campo rupestre, the negative effects of parasitic plants on host performance can be amplified, leading to a higher mortality rate of parasitized individuals 19, 14]. In fact, in this work, the individuals of the dominant species, *B. dracunculifolia*, that were parasitized showed a mortality rate of 61% after two years. In another work in the campo rupestre, of the 164 dead individuals of the species *M. calodendron*, 139 (84%) showed traces of the mistletoe *S. flexicaulis* [9].

The presence of *S. flexicaulis* altered the structure of the plant community in this study area, leading to decreased dominance and increased diversity and evenness of the plant community over time. This result was determined by the high mortality of parasitized individuals of the most abundant species. If the most parasitized individuals in the community belong to competitively dominant plant species, parasitism can mediate the coexistence of species in the community, allowing inferior competitor species to persist [11, 29, 31]. Similar effects of the presence of other aerial parasitic plants on the structure of plant communities have been demonstrated in different ecosystems [29, 51]. An experimental study in a coastal swamp environment demonstrated that the coexistence between plant species was mediated by the parasitism of the vine *Cuscuta salina*, since its preferred host plant is competitively dominant [29]. In another study, [51] demonstrated that the presence of the parasite *Cuscuta howelliana* led to an increase in richness and a change in the composition of species of plant communities that inhabit seasonal lakes. Similar results were also found for root hemiparasites [27, 30, 31]. Studies carried out with hemiparasitic species of the genus *Castilleja* in alpine ecosystems have shown that the presence of these species is associated with an increase in plant species richness and an increase in the evenness of the plants in the community [30, 31].

In this study, we determined the factors related to the characteristics of the host plants that affect the distribution and abundance of the mistletoe S. *flexicaulis,* and the effects of this parasitism on host plants and, consequently, on the structure of the communities they inhabit in the campo rupestre. The aggregate distribution of *B. dracunculifolia*, the dominant host in the community, seems to contribute to the parasite's success, since dense patches are more visible by birds in the landscape and facilitate the vegetative propagation of the parasite's individuals. In addition, we demonstrate that the architectural complexity of the host plant is an important factor that positively affects the abundance of the parasitic plant. In turn, the parasitism by S. *flexicaulis*, negatively affects the growth of host plants, which also have a higher mortality rate compared to non-parasitized plants. The high mortality of the most abundant species *B. dracunculifolia* led to a change in the structure of the plant community over time, with a decrease in dominance and an increase in the diversity and evenness of plant species. These results reinforce the importance of parasitic plants as key species for maintaining species diversity in plant communities.

## Acknowledgments

We thank two anonymous reviewers for their criticisms on earlier versions of the ms. We also thank the logistic support provided by the Reserva Vellozia and the Conselho Nacional de Desenvolvimento Científico e Tecnológico (CNPq) for funding this Long-Term Ecological Research (PELD-CRSC-17 - 441515/ 2016-9). We thank Maria Cristina Messias, Thaise Bahia and Hildeberto Caldas de Sousa for help in identifying plant species. We also thank Daniel Vasconcellos for translating this study to English. This study is in partial fulfillment of the requirements for a Doctor in Ecology at Universidade Federal de Minas Gerais (UFMG). CAPES provided scholarships to GFM, MB and SN. CNPq provided scholarships to YA and GWF.

